# Shifts in attentional demand are accompanied by sex-specific patterns of insula-frontal cortical interactions in mice

**DOI:** 10.64898/2025.12.02.691843

**Authors:** Abigail E. Harr, Matthew R. O’Leary, Eva Mei Vogt, Shreya Suresh, Henry L. Hallock

## Abstract

The insula is a brain region that is correlated with the cognitive process of attention in humans. Aberrant insula function is also implicated in attentional impairments in several psychiatric disorders, such as attention deficit hyperactivity disorder (ADHD), major depressive disorder (MDD), and schizophrenia. Although the insula’s role in attention has been well-studied in humans, we do not yet have an understanding of how the insula communicates with other brain areas to promote attentional processing. Here, we show that neurons in the mouse insula send direct axonal projections to the frontal cortex, and that these two brain areas functionally interact in different ways between male and female mice performing a touchscreen-based attention task. Specifically, we recorded and analyzed local field potentials (LFPs) from the insula and frontal cortex while mice performed variants of a rodent analogue of the continuous performance test of attention, or rCPT. In male mice, we found that activity in the frontal cortex preceded insular activity when the task was well-learned, but that insula-frontal cortex synchrony decreased when the need to inhibit responding was high. In female mice, synchrony between the insula and frontal cortex was largely bidirectional, but was high during all session types. We further demonstrate that patterns of theta-gamma coupling between the frontal cortex and insula distinguish between correct choices and errors during the rCPT, and that these patterns diverge between sexes. Our results suggest that functional differences in insula-frontal cortical circuitry at least partially underlie sex differences in attention-guided behavior in mice.

**Highlights:**

- We measured local field potentials (LFPs) in the mouse insula and frontal cortex during a translationally-relevant touchscreen-based task of attention
- Manipulation of stimulus frequency during the task increased the demand on vigilance and response inhibition
- In male mice, frontal cortex-driven synchrony in slow frequency bands reflected task rules, while theta-gamma coupling reflected errors when demand on response inhibition was increased
- In female mice, bidirectionally-driven synchrony in slow frequency bands also reflected task rules, while theta-gamma coupling reflected errors when demand on vigilance was increased

## 1. Introduction

Attention is the procedure by which material from the environment is selectively processed by the brain, allowing an organism to prioritize relevant sensory information (Knudsen, 2007). Attention involves multiple related, but distinct, processes, such as orienting, executive attention, and vigilance, which are controlled by multiple brain networks (Posner and Petersen, 1989; Petersen and Posner, 2012). Deficits in attentional function are implicated in many psychiatric disorders, such as attention deficit hyperactivity disorder (ADHD), and schizophrenia (Bush, 2010; Nuechterlein et al., 2015). It is therefore important to understand the neurobiological underpinnings of attention in order to identify potential diagnostic biomarkers, as well as therapeutic targets, for attentional symptoms in these disorders.

Much work using animal models in the past two decades has demonstrated that specific circuits (groups of interconnected neurons) in the brain have unique relationships to behavior and cognition (e.g., Namburi et al., 2015; Beyeler et al., 2016), including attention (Cope et al., 2019; Wang and Krauzlis, 2020; Norman et al., 2021). Results from these studies have led to the idea that cognitive symptoms in neuropsychiatric disorders, such as deficits in attention, could arise from aberrant function in these circuits (Gordon, 2016; Williams, 2016). Previous work from our lab characterized one such attentional circuit in mice - locus coeruleus (LC) neurons with axonal projections to the frontal cortex (FC; Hallock et al., 2024). We found that this circuit showed specific patterns of activation during a rodent analogue of a commonly used test of attention in humans: The continuous performance test, or CPT (Mar et al., 2013; Kim et al., 2015). During the CPT, participants must discriminate between a “target” stimulus, and a “non-target” stimulus. When the target stimulus is presented, participants must perform an action (such as pressing a button), and when the non-target stimulus is presented, participants must withhold responding (Rosvold et al., 1956; Conners et al., 2003). Specifically, our data showed that interactions (as measured through synchronicity of local field potential (LFP) recordings) between these brain areas in male mice was highest when the rodent CPT (rCPT) task was well-learned, providing potential electrophysiological markers of attentional function in this circuit.

Individual circuits, however, do not work in isolation; neurons in a given brain region form many circuits with other areas of the brain, including the FC and LC. One potential modulator of function in the LC-FC circuit is the insula, which is reciprocally connected with both brain areas in rodents (Gehrlach et al, 2020; Kayyal et al, 2021) and primates (Mesulam and Mufson, 1982) and is also implicated in a range of functions including, but not limited to, emotion, interoception, motivation, reward expectation, and attention (Corbetta et al., 2008; Centanni et al., 2021). Altered insula structure, function, and/or connectivity has also been observed in many psychiatric disorders, including bipolar disorder, schizophrenia, and ADHD (Ellard et al., 2018; Lopez-Larson et al., 2012; Sheffield et al., 2020; Shepherd et al., 2012). Insula abnormalities in psychiatric disorders are associated with a variety of cognitive impairments, including impairments in sustained attention (He et al., 2013; Lopez-Larson et al., 2012). For example, patients with bipolar disorder, as well as their relatives, display decreased accuracy on the CPT, and exhibit greater insula activation during error trials compared to healthy controls (Sepede et al., 2012). The presence of insula dysfunction in so many disorders highlights the need to understand this structure and its contribution to basic cognitive processes, including attention.

As a first step toward addressing this need, we first used a viral tracing approach to verify that the mouse insula sends projections to both the FC and LC. We then implanted depth electrodes to record LFPs from the FC and insula simultaneously while mice performed the rCPT. In addition to different brain regions, the current study also differs from the experiment performed in Hallock et al. (2024) in several important ways. First, we measured LFPs in both male and female mice in order to observe whether there are any sex-dependent differences in insula-FC connectivity during attention-guided behavior. Previous work has shown that male and female mice use different strategies during the rCPT (DeBrosse et al., 2023; Ramos et al., 2025; Li et al., 2025), suggesting that distinct neural processes drive sex-dependent behavior in this task. Second, we additionally measured LFPs during two probe sessions in the rCPT; in one probe session, the probability of encountering the target stimulus was decreased to 20%, and in the second probe session, the probability of encountering the target stimulus was increased to 80%. The rationale behind including these two extra probe sessions was to manipulate the demand on two specific attentional domains: Vigilance, and response inhibition. Previous research shows that errors of omission (withholding a response to the target stimulus) increase with decreased target stimulus probability (Beale et al., 1987; Denney et al., 2005), suggesting that this manipulation increases the demand for vigilance, or a sustained sense of alertness. Conversely, increasing target stimulus probability increases errors of commission (i.e., responding to the non-target stimulus; Wilson et al., 2016; Bedi et al., 2023; Mensen et al., 2024), suggesting that this manipulation increases the demand for response inhibition. In humans, the insula is involved in both vigilance (Fu et al, 2022) and response inhibition (Swick et al., 2011; Osada et al., 2024), but whether it accomplishes this by routing information about these distinct domains to different brain regions is not known. We found that, in male mice, insula-FC synchrony was largely driven by the FC, and distinguished between correct and error trials when target stimulus probability was high (higher demand on response inhibition). In contrast, insula-FC synchrony was either bidirectional or weakly driven by the insula in female mice, and distinguished between correct and error trials when target stimulus probability was low (higher demand on vigilance). Our results point to differences in insula-FC function as a potential contributor to sexually dimorphic performance in the rCPT.

## 2. Materials and Methods

### 2.1 Subjects

Subjects were 14 (7 male, 7 female) C57BL/6J mice (Jackson catalog # 000664). Mice were housed by sex, had free access to water, and were kept on a 12 hr light/dark cycle, with all behavioral experiments taking place during the light cycle. During behavioral experiments, mice were lightly food restricted to ∼5 grams of chow per day. Mice were between 3 - 5 months of age during testing. All procedures were in accordance with the Institutional Animal Care and Use Committee of Lafayette College.

### 2.2 Surgical Procedures

Two mice (1 male, 1 female) underwent surgery for the injection of a viral vector for retrograde labeling of afferents to the FC and LC. These viruses (AAVrg-Ef1a-EGFP; Addgene catalog # 55636-AAVrg, and AAVrg-CAG-tdTomato; Addgene catalog # 59462-AAVrg) contained plasmids coding for the expression of enhanced green fluorescent protein (EGFP) and tdTomato. For these surgeries, mice were first anesthetized with 2-3% isoflurane and the head was shaved. Mice were then placed in a stereotaxic frame (Kopf Instruments) with a nose cone administering a constant flow of isoflurane. Meloxicam was administered intraperitoneally as a systemic analgesic and lidocaine was administered subcutaneously above the skull as a topical analgesic. An incision was made along the scalp to expose the skull, and betadine was applied to sterilize the incision site. Two holes were drilled; one above the frontal cortex, +1.7 mm anterior to bregma, and ±0.3 mm lateral from the midline, and one above the LC, -5.4 mm posterior to bregma, and ±0.9 mm lateral from the midline. An automated infusion pump (World Precision Instruments, Sarasota, FL) was used to inject the virus at 4 nl/sec for a total volume of 600 nl unilaterally into the FC (1.7 mm ventral to the surface of the brain), and a total volume of 300 nl unilaterally into the LC (3.0 mm ventral to the surface of the brain). Following a six week incubation period, mice were transcardially perfused.

All other mice (12 total mice, 6 males, 6 females) underwent surgery to chronically implant depth electrodes (stereotrodes: Pinnacle catalog # 8425) for LFP recordings. For these surgeries, we attached a headstage containing four components: one electrode implanted in the frontal cortex, one electrode in the insula, one ground screw, and one reference screw. Mice were first anesthetized with 2-3% isoflurane and the head was shaved. Mice were then placed in a stereotaxic frame (Kopf Instruments) with a nose cone administering a constant flow of isoflurane. Meloxicam was administered intraperitoneally as a systemic analgesic and lidocaine was administered subcutaneously above the skull as a topical analgesic. An incision was made along the scalp to expose the skull, and betadine was applied to sterilize the incision site. Holes were drilled in four places (all coordinates relative to bregma): for the frontal cortex electrode, +1.7 mm AP and ±0.3 mm from the midline; for the insula electrode, +1.7 mm AP and ±2.5 mm from the midline. For the ground screw, a hole was drilled above the lambda skull suture, and for the reference screw, a hole was drilled in the left hemisphere (opposite from the insula and frontal cortex electrodes) over cortex. Electrodes were inserted into their respective locations and lowered to a depth of -1.7 mm for the frontal cortex electrode, and -1.9 mm for the insula electrode, relative to the surface of the brain. The headstage, electrodes, and wires were secured in place using dental cement (Lang Dental). Mice were given at least 5 days to recover before handling and behavioral testing.

### 2.3 Behavioral Training and Testing

Mice were trained on the rCPT according to methods previously described by our lab (Ramos et al., 2025). Specifically, mice were handled by the experimenter for ∼5 minutes per day for three to five days, and were exposed to the strawberry milkshake reward (a weighboat full of Ensure) in the homecage each day. Following handling, mice were habituated for two days by placing them inside the touchscreen chambers (Bussey-Saksida touchscreen chambers, Lafayette Instruments) for twenty minutes per day with a reward trough full of strawberry milkshake. Mice passed the habituation stage only if they consumed all of the reward by the end of the twenty minute period.

Following handling and habituation, mice were shaped to respond to a visual stimulus on the touchscreen at the front of the chamber. During this stage (stage 1), a white square was presented on the touchscreen. The square disappeared either when the mouse nosepoked it, or following a 10 s period. If the mouse nosepoked the square, a 3 kHz tone played, and a light illuminated the reward trough. When the mouse broke an infrared beam over the reward trough, the next trial was initiated, and another white square appeared on the touchscreen. Mice passed stage 1 when they reached a criterion of >40 nosepokes (“hits”) in a 30 minute session. Stage 2 was identical to stage 1, with the exception that mice were presented with a “go” stimulus (S+), which consisted of black and white horizontal or vertical bars (2 s stimulus duration, bar orientation counterbalanced across animals), instead of a white square.

During stage 3, mice were introduced to the no-go stimulus, or S- (a snowflake), and mice were shaped to withhold a response from this stimulus. There was a 50% chance that the mouse was presented with the S+ or S- during stage 3 sessions. There were four possible responses during stage 3 sessions: False alarms (the mouse incorrectly responds to the S-), correct rejections (the mouse correctly withholds response to the S-), hits (the mouse correctly responds to the S+), and misses (the mouse incorrectly withholds response to the S+). False alarms were punished with a 20 s timeout period. The time between stimulus presentations (intertrial interval or ITI) for stage 3 ranged from 2-3 s, and was random for each new trial.

To measure performance during stage 3 sessions, we calculated the mouse’s hit rate and false alarm rate. The hit rate is a metric of how frequently the mouse responded correctly to the S+, and was calculated by dividing the number of hit trials by the total number of S+ presentations. Similarly, the false alarm rate refers to how frequently the mouse incorrectly responds to the S-, and was calculated by dividing the number of false alarm trials by the total number of S- presentations:

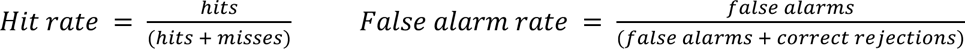

A mouse’s ability to discriminate between the S+ and S- and respond accordingly was quantified by the variable d’, which was calculated by subtracting the z-scored false alarm rate from the z-scored hit rate (Kim et al., 2015):

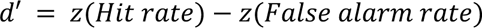

In order to pass stage 3, mice were required to achieve a d’ of at least 0.6 for two consecutive sessions. We additionally measured the mouse’s response bias (did the mouse have a tendency to respond more liberally or more conservatively) using a c score, with higher scores representing more conservative response bias, and lower scores indicating a more liberal response bias:

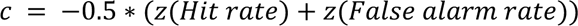

In order to examine how stimulus frequency affects insula-frontal cortical activity, we trained mice on two additional probe stages following stage 3 completion. In these stages, we manipulated the probability of S+ presentation. During probe 1, the S+ was presented roughly 20% of the time, and the S- 80% of the time. Decreasing the frequency of S+ presentations increases the need for vigilance, and errors of omission typically increase (Beale et al., 1987; Denney et al., 2005). During probe 2, the S+ was presented 80% of the time, and the S- 20% of the time. In contrast to probe 1, increasing the frequency of S+ presentations increases the need for response inhibition, and errors of commission typically increase (Wilson et al., 2016; Bedi et al., 2023; Mensen et al., 2024). Completion criteria for both probes was identical to stage 3 (d’ ≥ 0.6 for two consecutive days). The order in which mice were trained on probe 1 and probe 2 sessions was counterbalanced. In order to analyze neural activity during unique behavioral events, we extracted timestamps from our behavioral software (ABET; Lafayette Instruments) during hits and false alarms (timestamps were recorded when the mouse nosepoked the screen). Male and female mice were run in separate chambers for the duration of the experiment, and chambers were thoroughly cleaned with 50% ethanol between mice.

### 2.4 Electrophysiological Recordings

In order to assess the activity of neuronal populations within the insula and frontal cortex during learning, stimulus discrimination, and attention, simultaneous LFPs were recorded from these regions at various points during rCPT training and testing. LFP recordings were done at five timepoints throughout the experiment: On the last day of stage 2, on the first and last days of stage 3 (to see differences in activity when the task is not learned vs well-learned), and on the last days of both probe sessions (to see the effect of manipulating stimulus frequency on neural activity). A tether with a pre-amplifier was attached to the mouse’s headstage prior to each recording session. The tether was attached to a commutator at the top of the chamber, which fed into a data acquisition box that was also attached to the top of the chamber. LFPs were recorded using Sirenia Acquisition software (Pinnacle) with a sampling rate of 2 kHz, a 0.5 Hz high-pass filter, and a 200 Hz low-pass filter. In order to synchronize behavioral and electrophysiological data, we programmed the ABET software to send a TTL out at the start of the behavioral schedule, which was recorded as a timestamp in Sirenia. We used behavioral timestamps to isolate 4 s segments of LFP (2 s prior to, and 2 s following, screen touch) surrounding hits and false alarms during each recording session. LFP data was detrended in order to remove drifting artifacts (Bertocci et al., 2002), and trials with noise contamination (large amplitude 60 Hz oscillations) were identified and flagged using custom MATLAB functions. Trials were also visually inspected, and any trials that contained clipping artifacts (LFP signal that exceeded the y-axis threshold during recording) were also flagged. All flagged trials were excluded from further analysis.

### 2.5 LFP Analysis

To quantify the power (amplitude) of distinct oscillations within the insula and frontal cortex LFPs during rCPT behavior, we used the Chronux toolbox (Bokil et al., 2010) in MATLAB to perform multitaper power spectral density analysis for each trial. We further averaged power across two distinct frequency bands: The delta frequency band (0-4 Hz), and the theta frequency band (4-12 Hz), and compared averaged power in these frequency bands within-sex across response types (hits vs. false alarms) and sessions with mixed-factorial ANOVAs in R.

To assess the degree to which the insula and frontal cortical LFPs functionally interact during the rCPT, we also used the Chronux toolbox in MATLAB to calculate phase coherence, which measures the degree to which phases in defined frequency bands temporally synchronize over successive cycles of an oscillation. We again averaged coherence values across delta and theta frequencies, and compared these values within-sex between response types and sessions with mixed-factorial ANOVAs in R. To assess directionality between insula and frontal cortical LFPs, we performed Granger causality analysis with custom MATLAB functions, which determines the extent to which current continuously sampled signals from one brain area predict future continuously sampled signals from another brain area, above and beyond the extent to which signals from one brain area predict signals from the same brain area (Seth et al, 2015). This analysis produces two Granger causal indices: One index quantifies the extent to which signals in brain region “A” predict signals in brain region “B”, and the second index quantifies the extent to which signals in brain region “B” predict signals in brain region “A”. To turn these values into one metric, we took the Granger causal index for frontal cortex-to-insula directionality, and divided it by the total of both Granger causal index values to create a lead index. Lead index values above 0.5, therefore, suggest that frontal cortex directionality is predominant, while values below 0.5 indicate that insula directionality is more dominant. For all Granger causality analyses, we used the Akaike information criterion (AIC) to determine the optimal model order. We averaged lead index values across delta and theta frequency bands, and compared these values within-sex between response type and session with mixed-factorial ANOVAs in R.

Finally, we quantified the degree to which oscillations in distinct frequency bands interacted with each other by using custom MATLAB functions to assess phase-amplitude coupling between theta and gamma (50 - 120 Hz) oscillations in the frontal cortex and insula. To do this, we used a third degree Butterworth filter to extract theta and gamma oscillations from the raw LFP, identified peaks and troughs from the theta signal, and used this information to interpolate phases for each theta cycle. We then created a distribution of normalized (averaged) gamma power for each theta phase bin (18 bins between 0 and 360 degrees), and compared our observed theta-gamma distribution with a uniform distribution using Kullback Leibler divergence (Tort et al., 2010). This method generates a modulation index value - the higher this value, the higher the difference between the observed theta-gamma distribution and a uniform distribution, and the higher the phase-amplitude coupling. Modulation index values were statistically compared within-sex across response types and sessions using mixed-factorial ANOVAs in R.

### 2.6 Perfusions

Mice were transcardially perfused with 4% paraformaldehyde, brains were extracted and stored in 4% paraformaldehyde for 24 h, and then sunk in a 30% sucrose solution. For the two mice that received viral injections, coronal sections (50 μm) of the entire brain were cut on a sliding microtome (Leica) with an attached freezing stage (Physitemp). For the remaining mice, slices of the frontal cortex and insula were taken. Floating sections were incubated in a 1:5000 DAPI (Sigma) solution in 1x PBS for 20 m, and then mounted and coverslipped. Images were taken with an LSM 800 confocal microscope (Zeiss).

## 3. Results

### 3.1 Viral Tracing

To identify axonal projections to the FC and LC, we injected retrograde viruses encoding EGFP and tdTomato, respectively, into these areas of the brain (Fig. 1a), and took whole brain slices to look for EGFP and tdTomato expression. The insula exhibited a strong fluorescent signal (Fig. 1b), which is in line with previous research demonstrating direct reciprocal connections between the insula, FC, and LC (Gehrlach et al, 2020). We chose to focus on connections between the FC and insula for subsequent experiments.

**Figure 1:**
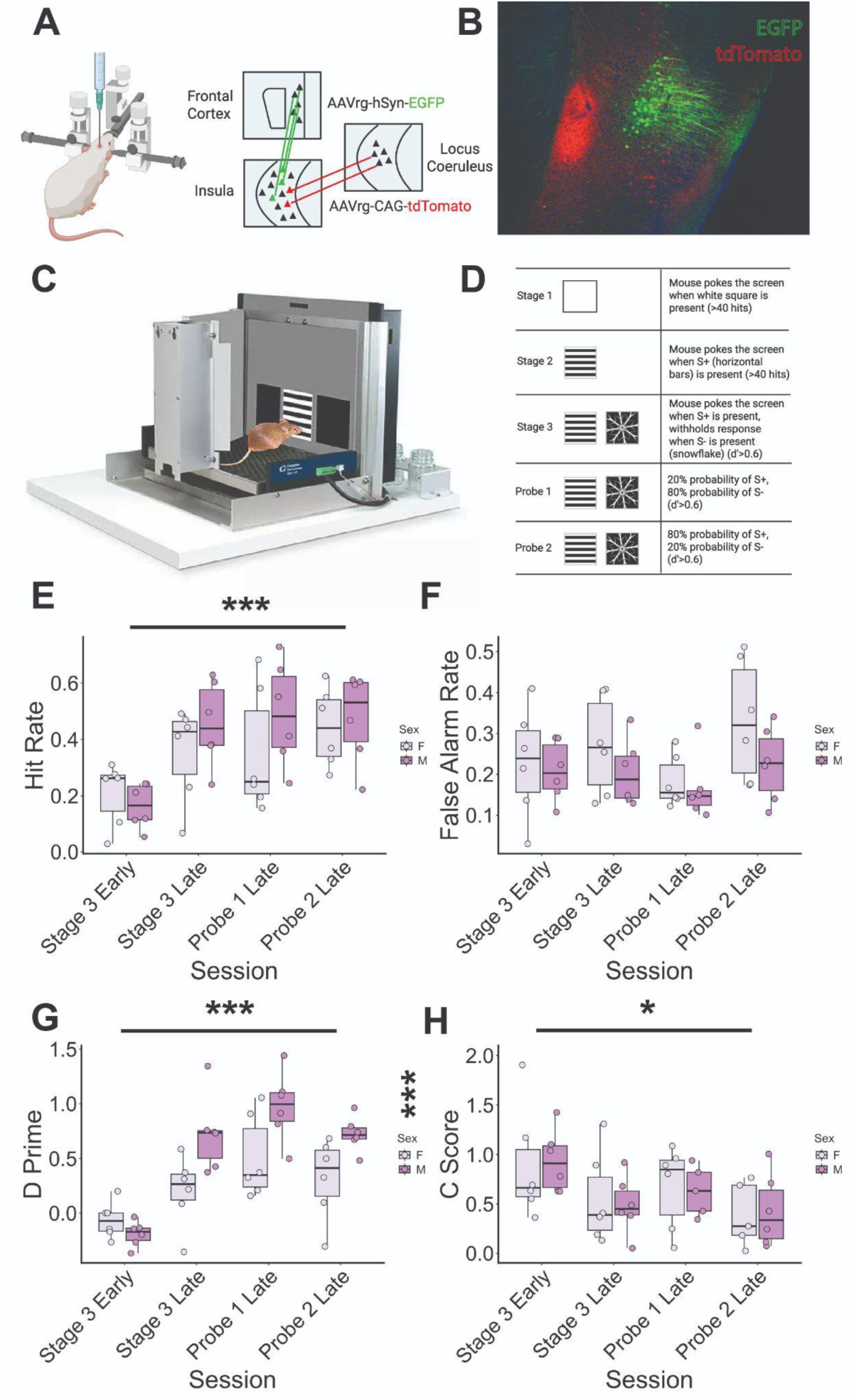
Insula-FC connectivity and behavior during rCPT sessions. *(A)* Schematic of injection strategy for viral labeling of insula neurons with projections to the FC. *(B)* Coronal section of the insula showing eGFP expression from a retrograde virus injected into the FC, and tdTomato expression from a retrograde virus injected into the LC. *(C)* Cartoon depiction of the touchscreen chamber used for the rCPT, with touchscreen at the front, and reward trough in the back. *(D)* Descriptions of rCPT stages and behavioral criteria for passing onto the next stage. *(E)* Hit rate between male and female mice across rCPT sessions. Hit rate significantly differed across sessions in both sexes. *(F)* False alarm rate between male and female mice across rCPT sessions. No significant differences in false alarm rate were observed as a function of session or sex. *(G)* D’ (the ability of mice to discriminate between the target and non-target) significantly increased after stage 3 early sessions in both sexes, but d’ was significantly lower in female mice across sessions. *(H)* C score (liberal vs. conservative response strategies) was significantly lower after stage 3 early sessions in both sexes. * = *p<*0.05, *** = *p*<0.001, sex x session ANOVAs.

### 3.2 rCPT Behavior

In order to examine how the frontal cortex and insula interact during attention-guided behavior, we implanted depth electrodes into these brain areas, and recorded LFPs during distinct rCPT stages (see Fig. 1c for chamber schematic, Fig. 1d for description of stages). When comparing across these stages, the animals’ hit rate significantly changed across sessions (*F*(1,5) = 4.355, *p* = 0.0019, main effect of session), increasing after the task was well-learned (from stage 3 early to stage 3 late), and then leveling off (Fig. 1e). Between-sex differences did not reach statistical significance (*p* = 0.0605), but male mice’s hit rates were generally higher than that of female mice. In contrast, false alarm rates did not significantly differ as a function of either sex or session, but were higher during probe 2 sessions compared to probe 1 sessions in both sexes (Fig. 1f), possibly reflecting the tendency for errors of commission to increase when stimulus frequencies also increase (Wilson et al., 2016; Bedi et al., 2023; Mensen et al., 2024).

When assessing d’ scores, which measure an animal’s ability to discriminate between an S+ and S-, we found significant main effects of session (*F*(1,5) = 9.795, *p* < 0.001), and sex (*F*(1,5) = 17.223, *p* < 0.001). D’ scores were lowest during stage 3 early sessions in both sexes, and were significantly lower across all sessions in female mice (Fig. 1g). Correspondingly, c scores were highest during stage 3 early compared to other sessions in both sexes (*F*(1,5) = 2.594, *p* = 0.0344), indicating that mice responded more conservatively (nosepoked the touchscreen less) when the task rule was unknown (Fig. 1h).

### 3.3 LFP Recordings

To examine the contributions of the frontal cortex and insula to rCPT task performance, we first identified timestamps for hits and false alarms, and isolated 4 second segments of LFP (extending from 2 seconds previous to, and 2 seconds following, screen touch) from each trial in both brain regions (see Fig. 2a and Fig. 2b). We chose to focus on hits and false alarms because the mouse’s behavior is similar during both response types, whereas behavior during correct rejections and misses is more variable and not easily quantified. We verified that our electrodes targeted the FC and insula (Fig. 2c), and then analyzed relationships to behavior within each brain region by comparing power spectral density between response types and sessions. In both the insula and frontal cortex, we found that the amplitude of slow oscillations - namely delta (0-4 Hz) and theta (4-12 Hz) - dominated the LFP signal (Fig. 2d). Because of this, we chose to focus on these two frequency bands for our comparisons. We found that neither delta, nor theta, power changed significantly as a function of either response type or session in either brain region, for either sex (Fig. 2e, Fig 2f). This was likely due, in part, to the large amount of variability in power values in our sample, as mean power values were skewed by outliers in several sessions.

**Figure 2:**
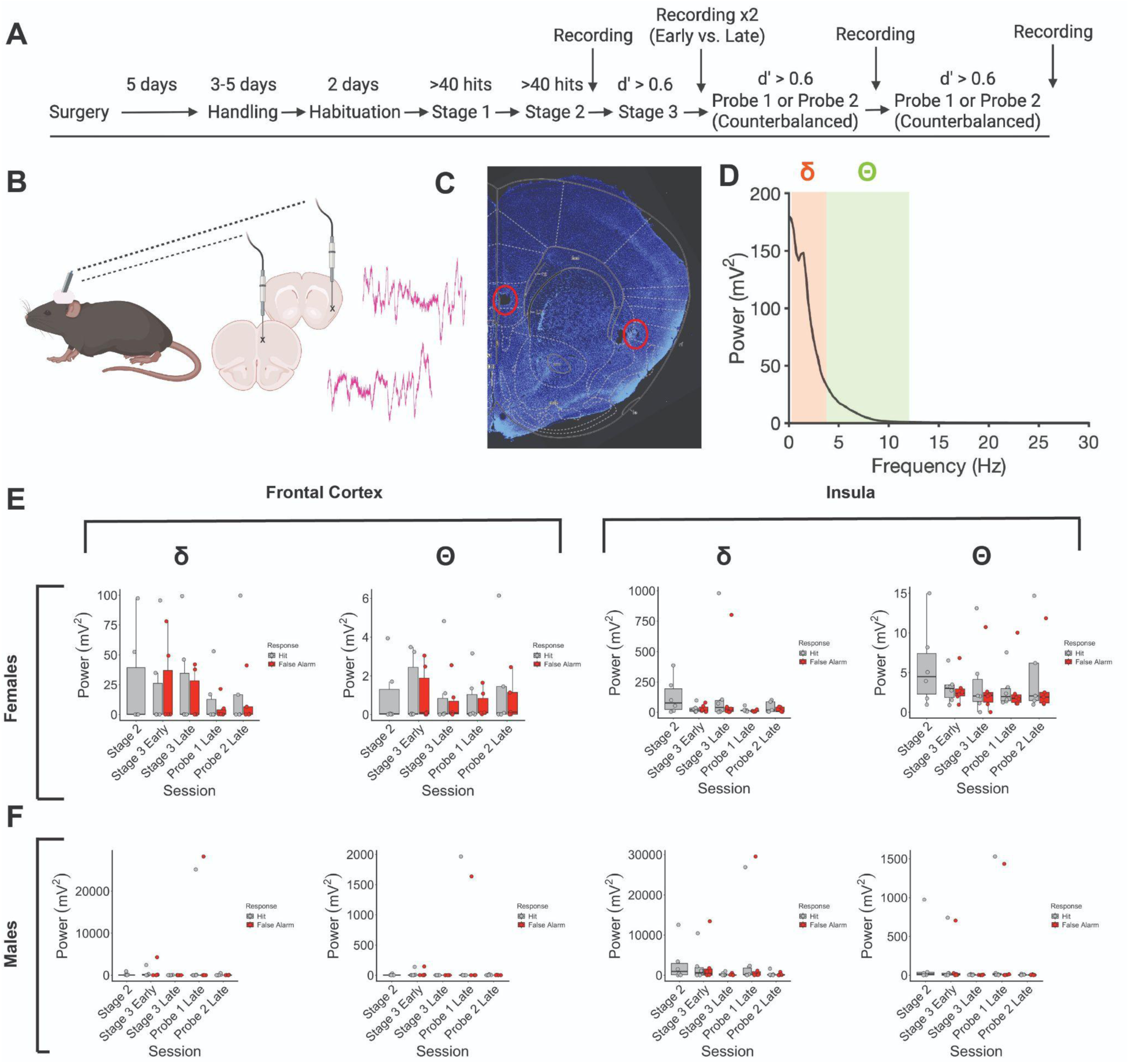
Power spectral density in the insula and FC during rCPT sessions. *(A)* Experimental timeline for rCPT training/testing and LFP recordings. *(B)* Schematic showing electrode placements in the insula and FC, with representative raw LFP traces from each brain region. *(C)* Representative coronal section of the FC and insula, with electrode tips inside of red circles. Blue staining = DAPI. *(D)* Example power spectral density plot, with delta (0-4 Hz) and theta (5-12 Hz) frequency bands highlighted in orange and green, respectively. *(E)* Neither power in the delta frequency band, nor power in the theta frequency band, differed significantly between sessions in females, *(F)* or males.

We next examined the degree to which the frontal cortex and insula interacted during the rCPT by measuring phase coherence in the two LFPs, defined as the degree to which within-frequency phases temporally align on successive cycles (Fig. 3a). In females, delta coherence was lowest during stage 2, and significantly increased during other sessions (*F*(1,4) = 2.758, *p* = 0.0397, main effect of session), indicating that the insula and prefrontal cortex functionally synchronize when female mice must discriminate between stimuli (Fig. 3b, Fig. 3c). Theta coherence in females followed a similar pattern, although no effects were statistically significant. Coherence in male mice was more complex; phase coherence in both delta and theta frequency bands changed significantly as a function of session (*F*(1,4) = 6.301, *p* < 0.001 for delta, *F*(1,4) = 6.9743, *p* < 0.001 for theta), increasing after stage 2, but decreasing again during probe 2 late sessions (when S+ frequencies were highest; Fig. 2b, Fig. 2d). This difference in phase coherence patterns between male and female mice suggests that interactions between the insula and frontal cortex reflect different information between sexes, signaling the need for stimulus discrimination regardless of S+ frequency in females, while signaling stimulus discrimination only when the demand on response inhibition is relatively low in males.

**Figure 3:**
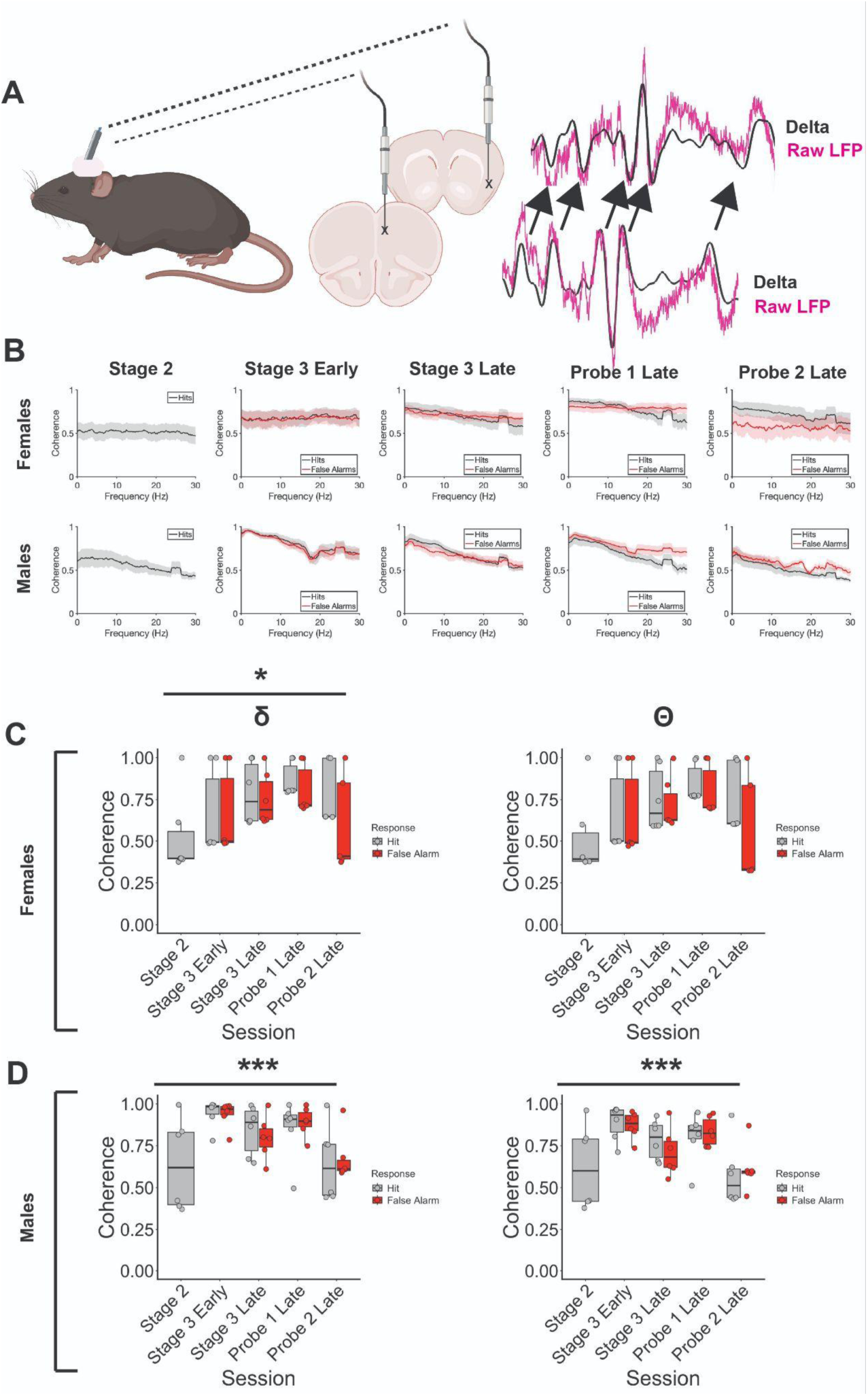
Phase coherence between delta and theta oscillations in the insula and FC during rCPT performance. *(A)* Schematic showing example insula and FC LFP traces in purple, with filtered delta oscillations overlaid in black. Arrows demonstrate phase consistency across delta cycles. *(B)* Plots showing phase coherence across frequency bands in females and males during different rCPT sessions. *(C)* Delta and theta coherence across sessions and response types in female mice. Delta coherence significantly increased after stage 2 sessions, and was generally higher during hits vs. false alarms. *(D)* In male mice, delta (left panel) and theta (right panel) coherence increased significantly after stage 2 sessions, but decreased again during probe 2 sessions. * = *p*<0.05, *** = *p*<0.001. Shaded portions in *(B)* = SEM.

To determine directionality of information flow between the insula and frontal cortex, we used Granger causality to quantify the degree to which the LFP in either brain region influenced the LFP in the other brain region. We used this information to calculate a lead index, with values greater than 0.5 indicating that directionality is strongest from frontal cortex to insula, and values lower than 0.5 indicating that directionality is strongest from insula to frontal cortex (see Methods). In female mice, the insula signal strongly predicted the frontal cortex signal in delta and theta frequency bands during stage 2, but this effect got weaker as mice learned the task and progressed through the probe sessions (Fig. 4a, Fig. 4b). There were no differences in directionality observed in any frequency band between response types (hits vs. false alarms) in female mice. Taken together with the results of our phase coherence analysis, these data suggest that the flow of information during learning in the rCPT sessions shifts from insula-driven to bidirectionally-driven, and that this shift is accompanied by a corresponding increase in synchrony between the two brain areas, possibly reflecting learning and implementation of the general task rule. We observed a different pattern in male mice, such that frontal cortex-to-insula directionality dominated during all well-learned sessions (all sessions with the exceptions of stage 2 and stage 3 early; Fig. 4a), especially in the theta frequency band (*F*(1,4) = 4.519, *p* = 0.00374, main effect of session; Fig. 4c). Similarly to females, directionality did not differ between response types in males. Taken together with our phase coherence results in males, our data indicate that, in contrast to females, the frontal cortex strongly influences the insula during rCPT performance, and that this connection is only weakened during probe 2, when the need for response inhibition is increased.

**Figure 4:**
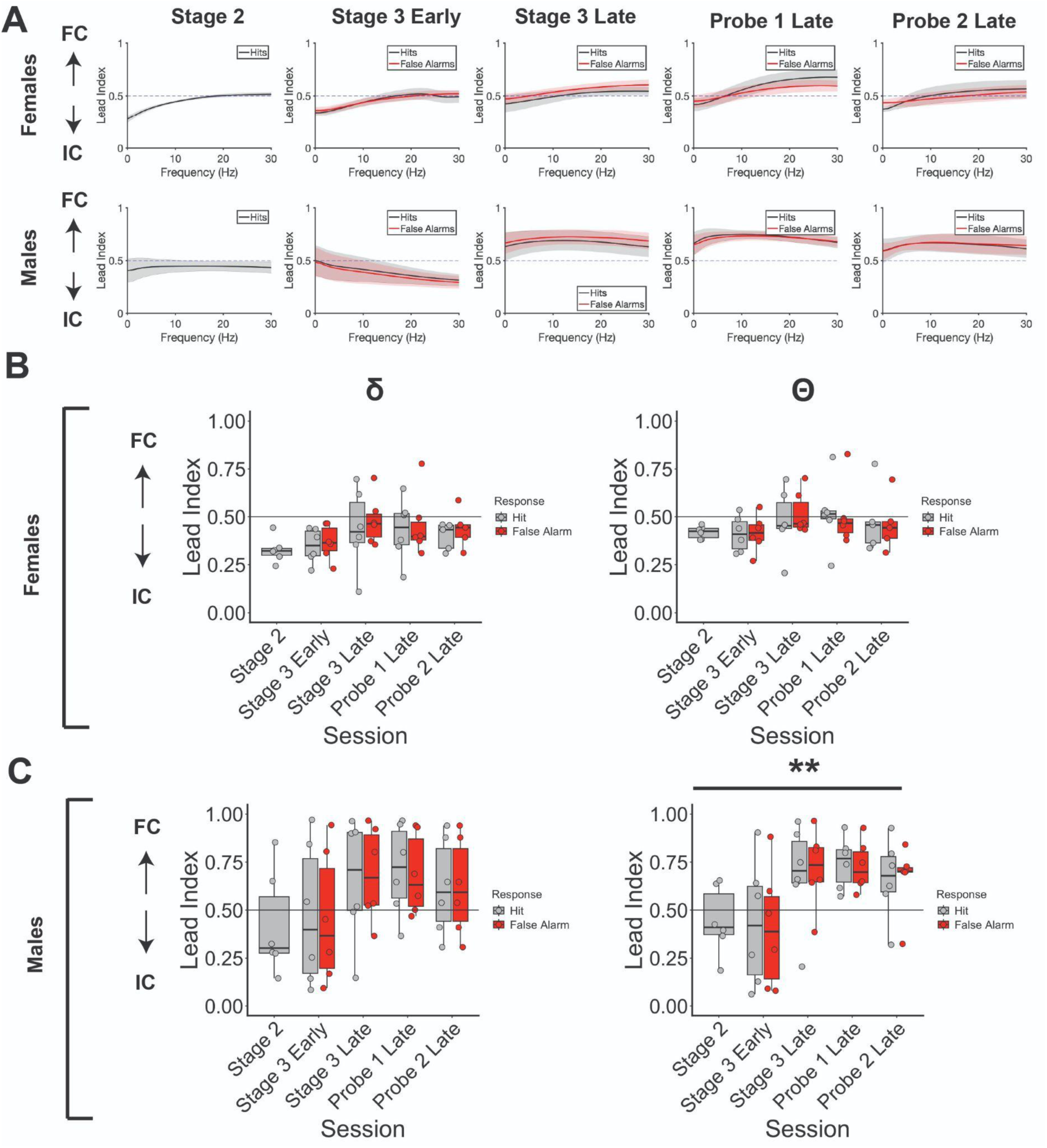
Directionality between insula and FC LFPs during the rCPT. *(A)* Granger causality demonstrates differences in which brain region leads the other during distinct rCPT sessions in male and female mice. In female mice (top row), the insula leads the FC (values less than 0.5 on the y-axis) in delta and theta frequency bands during stage 2 and stage 3 early sessions, but this effect decreases in later sessions. In male mice (bottom row), the FC strongly leads the insula (values greater than 0.5 on the y-axis) during all well-learned rCPT sessions. *(B)* No significant differences in lead index between sessions or response types were observed in delta or theta frequency bands in female mice. *(C)* In male mice, lead index in the theta frequency band significantly increased after stage 3 early sessions. ** = *p*<0.01, shaded portions in *(A)* = SEM.

Oscillations in different frequency bands can also communicate with one another, and this phase-amplitude coupling correlates with behavioral performance and cognition (Bazzigaluppi et al., 2018; Radiske et al., 2020), including during tests of attention (Szczepanski et al., 2014; Chacko et al., 2018). Theta-gamma coupling (the tendency for bouts of high-amplitude gamma (40-100 Hz) to occur on the same phase of a theta cycle across time) in particular has a strong relationship with cognition and behavior (Tamura et al., 2017; Barr et al., 2017; Koster et al., 2018). To determine whether theta-gamma coupling supports performance in this task, we measured theta-gamma phase-amplitude coupling both between, and within, the frontal cortex and insula during the rCPT (Fig. 5a). We found strong evidence for theta-gamma coupling (as opposed to between other frequency bands), particularly in gamma frequency ranges between 80 and 120 Hz (Fig. 5b), with the ascending phase (0-180 degrees) of theta modulating the amplitude of these gamma oscillations (Fig. 5c). In female mice, frontal cortex-to-insula theta-gamma coupling distinguished between response types only during certain sessions, being higher during hits during stage 3 late, and probe 2 sessions (*F*(1,4) = 5.129, *p* = 0.00181, main effect of session, *F*(1,3) = 3.934, *p* = 0.01445, response type x session interaction; Fig. 5d). In contrast, insula-to-frontal cortex theta-gamma coupling signaled errors (false alarms) during probe 1 sessions when S+ frequency was low (*F*(1,4) = 4.909, *p* = 0.00507, response type x session interaction; Fig. 5d). Theta-gamma coupling within the frontal cortex was similar, increasing during probe 1 false alarms (*F*(1,4) = 3.704, *p* = 0.0186, response type x session interaction; Fig. 5d). Theta-gamma coupling in the insula of female mice generally decreased between stage 2 and other sessions (*F*(1,4) = 3.235, *p* = 0.0209, main effect of session), but did not distinguish between response types.

**Figure 5:**
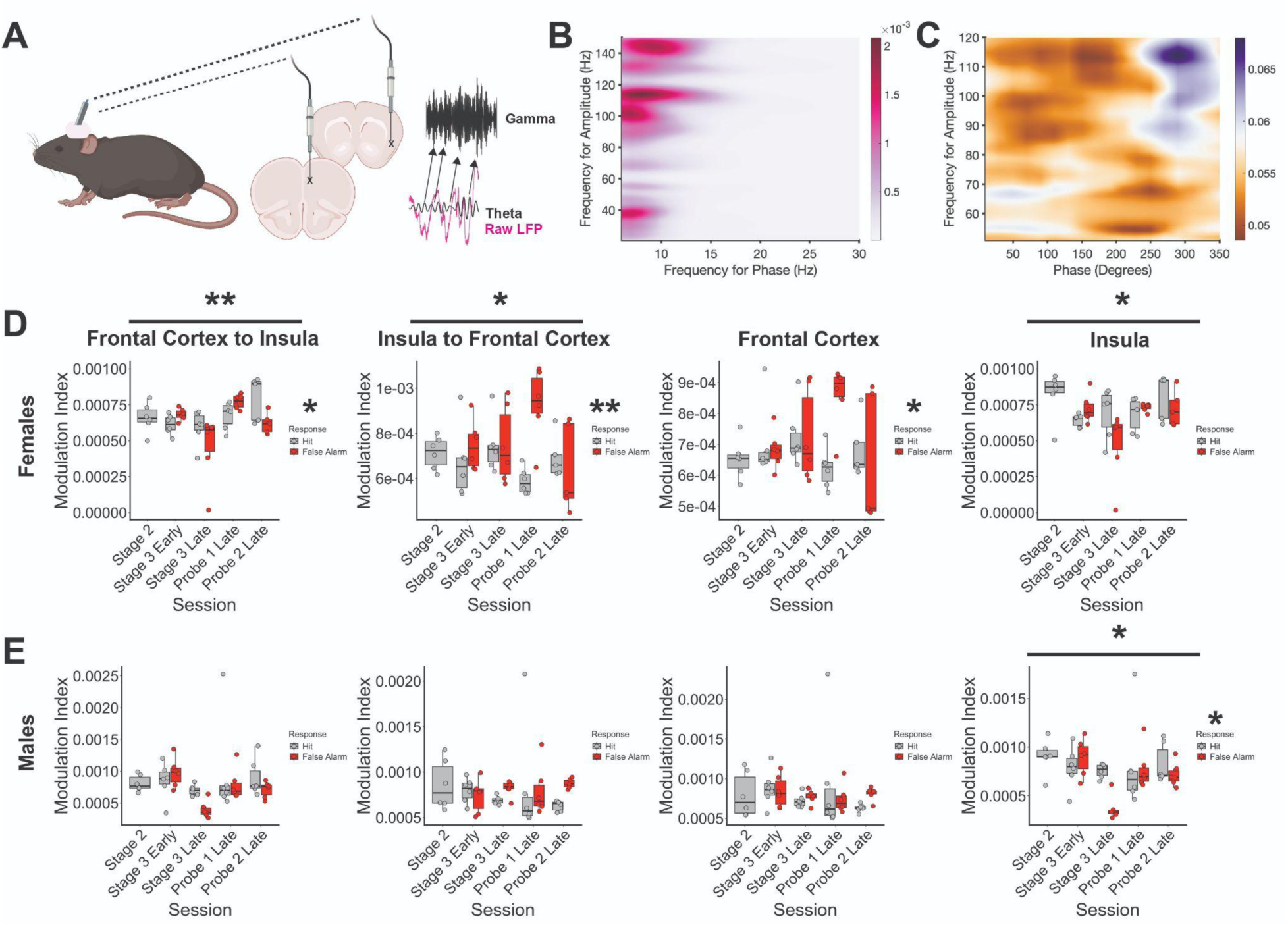
Theta-gamma phase-amplitude coupling in the insula and FC during the rCPT. *(A)* Schematic of coupling between theta phase in the frontal cortex (bottom LFP trace) and gamma amplitude in the insula LFP (top trace). Arrows link similar theta phases to bouts of high amplitude gamma. *(B)* Co-modulogram demonstrating consistent phase-amplitude relationships between delta/theta oscillations (x-axis) and mainly fast gamma oscillations (>100 Hz, y-axis) in the FC from a representative mouse. Color = modulation index. *(C)* Phase map showing higher modulation of fast gamma amplitude (purple region) during ascending phases of theta (theta trough = 180 degrees, theta peak = 360 degrees on the x-axis) from the FC of a representative mouse. Color = modulation index. *(D)* FC theta-to-insula gamma phase-amplitude coupling significantly differed as a function of session in females, with a session-by-response type interaction also being observed (first panel). Both insula (second panel) and frontal cortex (third panel) theta-to-frontal cortex gamma coupling significantly differed between response types, and insula theta-to-insula gamma coupling significantly differed across session types (fourth panel). *(E)* In male mice, frontal cortex theta-to-insula gamma coupling significantly differed between response types (first panel), and insula theta-to-insula gamma coupling significantly differed across sessions (fourth panel). * = *p*<0.05, ** = *p*<0.01.

In male mice, insula-to-frontal cortex theta-gamma coupling was higher during false alarms during stage 3 late and probe 2 sessions, but these differences did not survive multiple comparisons (Fig. 5e). Similarly, theta-gamma coupling within the frontal cortex did not significantly differ as a function of session or response type, even though it was higher during stage 3 late and probe 2 sessions. Theta-gamma coupling within the insula changed across sessions (*F*(1,4) = 2.979, *p* = 0.0289, main effect of session), and was significantly lower during stage 3 late false alarms (*F*(1,1) = 4.319, *p* = 0.0434, main effect of response type; Fig 5e).

These effects reveal multiplexed signals within the insula-frontal cortex circuit that differ between sex, session type, and response type. In female mice, bidirectionally-driven synchrony between low-frequency oscillations was higher during all sessions that required stimulus discrimination, while theta-gamma coupling from the insula-to-frontal cortex and within the frontal cortex was higher during probe 1 false alarms (when the demand on vigilance is increased). In male mice, frontal cortex-driven synchrony between low-frequency oscillations did not broadly distinguish between response types, but did distinguish between sessions (decreasing during probe 2 sessions), while insula-to-frontal cortex and frontal cortex theta-gamma coupling was higher during probe 2 false alarms.

## 4. Discussion

In the current study, we confirm previous findings that the mouse insula sends direct axonal projections to the frontal cortex, or FC (Gehrlach et al, 2020; Kayyal et al, 2021), and find that these two brain areas functionally communicate during the performance of a translationally-relevant touchscreen-based attention task (the rCPT). Specifically, we demonstrate that, in male mice, slow oscillations in the insula and FC LFPs synchronize when the task rule is well-learned, and this synchrony is strongly driven by the FC. However, phase synchrony decreases in males when demand on response inhibition increases (i.e., when target frequency increases), and theta-gamma phase-amplitude coupling between the insula and FC increases specifically during false alarms in these sessions. These results suggest that, in males, the FC drives communication with the insula to promote the use of learned task rules; however, when target probability increases, communication in this circuit switches to a different frequency, and this communication increases during errors of commission. This result intimates that the insula-FC circuit encodes impulsive responding in males in situations where the need to suppress impulsivity is increased.

Similarly to male mice, functional synchrony between slow oscillations in the insula and FC LFPs of female mice was generally higher across all sessions that required stimulus discrimination, possibly reflecting general increases in task rule learning and utilization. In contrast to male mice, this synchrony remained relatively high during probe 2 sessions. Also in contrast to males, this synchrony was either bidirectional or weakly driven by the insula. Theta-gamma phase-amplitude coupling between the insula and FC was also higher during false alarms in females, but only when vigilance demand increased during probe 1 sessions (i.e., when target frequency was reduced). These results suggest that this circuit can encode both task rules and decreases in vigilance by using different frequency bands, and that how activity in this circuit responds to vigilance is dependent on the attentional context. These findings also highlight major sex differences in how the insula-FC pathway responds during attention-guided behavior: In male mice, a strongly FC-driven signal reflects task rules, but switches to encoding impulsivity in certain contexts, and in female mice, a bidirectionally-driven signal reflects both task rules and decreases in attention that are dependent on vigilance demand.

Previous research has shown that there exist gender differences in CPT performance in human adolescents with ADHD, with males exhibiting greater impulsivity (more errors of commission) than females (Hasson and Fine, 2012), which aligns with a constellation of findings that point to higher impulsivity in males generally (Chapple and Johnson, 2007; Cross et al., 2011; Weinstein and Dannon, 2015). Our lab recently demonstrated that male mice respond more liberally than female mice during initial rCPT training sessions (Ramos et al., 2025), possibly reflecting increased impulsive action. Another interpretation of this result, however, is that male mice are simply quicker to develop a task strategy (van den Bos et al., 2012; van den Bos and McClure, 2013), possibly because they are less risk-averse than female mice (Orsini et al., 2016). In the current study, both male and female mice had higher false alarm rates, and lower c scores, during probe 2 performance compared to probe 1 performance (Fig. 1f), supporting the hypothesis that increased target stimulus probability increases demand on response inhibition; however, probe 2 false alarm rates did not significantly differ between sexes, dovetailing with experiments that failed to find sex differences in impulsive action in rodent models (Anker et al., 2008; Papaleo et al., 2012; Burton and Fletcher, 2012).

Despite the lack of sex differences in impulsive-like behavior, male and female mice displayed drastically different patterns of insula-FC connectivity during probe 2 sessions. FC-driven communication in slow frequency bands was reduced during probe 2 sessions in males, and insula-FC theta-gamma coupling increased during errors of commission (false alarms). Research in human males shows increased anterior insula activation during inhibitory failure (increased impulsivity; Dambacher et al., 2015), while the role of the insula in response inhibition in females is mixed, with some studies showing increased insula activation in mixed-gender studies (Ramautar et al., 2006; Bastin et al., 2016), and some showing decreased insula activation in females with borderline personality disorder (BPD; Mortensen et al., 2016). Insula responsivity to impulsive choice may therefore be sex-dependent in both humans and rodents, with a greater role for the insula in failure to inhibit responding in males. The concurrent reduction in FC-driven delta and theta synchrony between the insula and FC during probe 2 sessions in males could reflect a simultaneous shift away from FC networks when demand on response inhibition increases. The rodent FC is most anatomically and functionally homologous with the human/non-human primate cingulate cortex (CC; Laubach et al., 2018; van Heukelum et al., 2020), and the anterior cingulate cortex (aCC) in humans functionally synchronizes with the parietal cortex during impulsive action (Golchert et al., 2017), while aCC-insula synchrony is more heavily associated with errors of omission (withholding a response; Jung et al., 2014). Taken together, these findings implicate shifts in communication between anatomically-distinct FC circuits during attention in males, with higher FC-to-insula communication during periods of low need for response inhibition, and a higher insula activation (possibly accompanied by shifts to other FC-driven circuits) when the need for response inhibition increases.

In contrast to probe 2 sessions, false alarm rate in probe 1 sessions (20% target frequency) was reduced in both male and female mice, which is in line with research demonstrating that reduced target frequency results in higher errors of omission (Beale et al., 1987; Denney et al., 2005). In females, theta-gamma coupling in the insula-FC circuit was higher during false alarms compared to hits, while insula-FC theta-gamma coupling did not distinguish between response type during probe 1 sessions in males. The fact that theta-gamma coupling in this circuit reflects errors of commission in both males and females suggests that insula-FC communication is important for error detection during attention tasks, which is in line with a large body of evidence in humans (Carter et al., 1998; Holroyd et al., 2004; Ullsperger et al., 2010; Harsay et al., 2018). However, our results show that whether this circuit responds to errors of commission is highly dependent on attentional context and sex. Male mice tend to respond more liberally (more likely to nosepoke the screen during any given trial) than female mice in the rCPT (Ramos et al., 2025), suggesting that probe 2 sessions may be more difficult for them because they have less baseline response inhibition. In contrast, female mice respond more conservatively than males, indicating that probe 1 sessions may be more difficult for them because they have higher levels of baseline response inhibition. If this is the case, then insula-FC theta-gamma coupling may be responsible for error detection during instances of increased attentional effort, which might shift depending on the strategy used by an animal to solve the task.

Human studies have shown sex differences in regional blood flow during attention tasks, including in the insula and ACC (Bell et al., 2006; Dumais et al., 2018; Gaillard et al., 2020), including studies that have shown greater ACC activation during impulsive responding in males (Liu et al., 2012). These differences in patterns of activation between males and females may be partially due to differences in catecholaminergic function between sexes. Catecholamines, such as dopamine and norepinephrine, heavily influence function in cortical circuits that control attention (Robbins, 1984; Puumala and Sirvio, 1998; Bymaster et al., 2002; Aalto et al., 2005; Dang et al., 2012), and sex differences in locus coeruleus (the major manufacturer of norepinephrine in the brain) and ventral tegmental area (one of the major manufacturers of dopamine in the brain) function may influence task strategy and cortical activity during tasks such as the rCPT (Curtis et al., 2006; Trainor, 2011; Bangasser and Valentino, 2012; Bangasser et al., 2016; Zachry et al., 2021). Catecholaminergic signaling is also heavily implicated in disorders of attention, such as attention-deficit hyperactivity disorder (ADHD; Biederman and Spencer, 1999; Swanson et al., 2000; Volkow et al., 2009; Wu et al., 2012; Vanicek et al., 2014), and sex differences in dopamine and norepinephrine pathways may partially underlie dissimilarities in how ADHD symptoms manifest in males and females (Andersen and Teicher, 2000; Fedele et al., 2012; Arnett et al., 2015; Mowlem et al., 2019). If this is the case, identifying sex-specific biomarkers of function in downstream attention circuits, such as the insula and ACC, could provide valuable diagnostic information for disorders that feature sex-specific deficits in attention, such as ADHD.

## Acknowledgements

The authors gratefully acknowledge Amy Badillo for animal care assistance.

## CRediT Author Contributions

**AEH**: Conceptualization, data curation, formal analysis, investigation, methodology, visualization, writing; **MRO:** Data curation, investigation; **EMV:** Data curation, investigation; **SS**: Data curation, investigation; **HLH**: Conceptualization, formal analysis, funding acquisition, project administration, resources, supervision, visualization, writing.

